# Interaction of NPC2 protein with Lysobisphosphatidic Acid is required for normal endolysosomal cholesterol trafficking

**DOI:** 10.1101/559849

**Authors:** Leslie A. McCauliff, Annette Langan, Ran Li, Olga Ilnytska, Debosreeta Bose, Peter C. Kahn, Judith Storch

## Abstract

Unesterified cholesterol accumulation in the late endosomal/lysosomal (LE/LY) compartment is the cellular hallmark of Niemann-Pick C (NPC) disease, caused by defects in the genes encoding NPC1 or NPC2. We previously reported the dramatic stimulation of NPC2 cholesterol transport rates by the LE/LY phospholipid lysobisphosphatidic acid (LBPA) and in these studies sought to determine their functional relationship in normal LE/LY cholesterol egress. Here we demonstrate that NPC2 interacts directly with LBPA and identify the NPC2 hydrophobic knob domain as the site of interaction. Using its precursor phosphatidylglycerol (PG), we show that PG-induced LBPA enrichment results in clearance of accumulated cholesterol from NPC1-deficient cells but is ineffective in cells lacking functional NPC2. Together these studies reveal a heretofore unknown aspect of intracellular cholesterol trafficking, in which NPC2 and LBPA function together in an obligate step of sterol egress from the LE/LY compartment, which appears to be independent of NPC1.

## Introduction

Cholesterol is a small, hydrophobic molecule that is a vital building block of cell membranes and a precursor for steroid hormones, bile salts, vitamin D, and oxysterol ligands for transcription factors. Intracellular transport of cholesterol is a highly regulated but, as yet, incompletely understood process. Perturbations can lead to detrimental outcomes such as in the lysosomal storage disorder Niemann Pick Type C (NPC) disease, where LDL-derived cholesterol becomes trapped within the late endosomal/lysosomal (LE/LY) system. The sterol particularly enriches LE/LY inner membranes, which develop during endosome maturation as a means of compartmentalizing its contents (Gruenberg, 2001; Gruenberg, 2003; Matsuo et al., 2004). The accumulation of cholesterol in NPC disease is associated with amassing of other lipids in the LE/LY, disruption of post-lysosomal cholesterol metabolism, and ultimately clinical manifestations including organomegaly and neurological deterioration. In 95% of NPC cases, mutations in the large LE/LY transmembrane protein, NPC1, prevent proper export of cholesterol from the LE/LY to other cellular compartments. The remaining 5% of cases are caused by mutations in the small, 132-amino acid, soluble LE/LY protein, NPC2 (Peake & Vance, 2010; Sokol et al., 2010).

Similarities in the cellular and clinical phenotypes resulting from either NPC1 or NPC2 deficiency have led to the suggestion that these two proteins function cooperatively in normal LE/LY cholesterol trafficking (Kwon et al., 2009; Sleat et al., 2004); a proposed model shows cholesterol directly transferred from NPC2, in the LE/LY lumen, to NPC1, located in the limiting membrane of the compartment (Estiu et al., 2013; Wang et al., 2010). Recent tertiary structural analyses identifying a potential NPC2 interacting domain on the NPC1 protein, support this mode of cholesterol egress from the LE/LY compartment (Zhao et al., 2016). It is further proposed that cholesterol transfer from NPC2 to the luminally localized N-terminal domain of NPC1 allows sterol passage through the glycocalyx found at the luminal surface of the LE/LY limiting membrane, via concerted effort from membrane glycoproteins (Li et al., 2016).

In addition to functioning with NPC1, accumulating evidence suggests that NPC2 may also have NPC1-independent actions. Goldman and Krise, for instance, showed that NPC2 deficient fibroblasts exhibit significant reductions in exocytosed dextran relative to NPC1 deficient cells. They additionally demonstrated that treatment with the commonly used inducer of the NPC disease phenotype, U18666A, further reduced exocytosis in NPC1 but not NPC2 cells, indicating a potential divergence in the egress pathway of membrane-impermeable species utilized by the NPC1 and NPC2 proteins (Goldman & Krise, 2010). It has also been shown that overexpression of ABCA1 reversed lysosomal cholesterol accumulation in NPC1 deficient but not NPC2 deficient cells (Boadu et al., 2012). Further, Karten and colleagues demonstrated that endosome to mitochondrial cholesterol transport occurs efficiently in the absence of NPC1 protein but is dependent upon the presence of functional NPC2 (Kennedy et al., 2012).

NPC2 binds cholesterol with a 1:1 stoichiometry (Xu et al., 2007). In 2003, Ko et al. showed that point mutations preventing cholesterol binding also prevented NPC2-mediated LE/LY cholesterol efflux from NPC2 deficient patient fibroblasts; cholesterol binding was therefore deemed essential to the function of NPC2 (Ko et al., 2003). Interestingly, however, some NPC2 point mutants with normal cholesterol binding affinity were nevertheless deficient in clearing intracellular cholesterol from NPC2 patient fibroblasts, suggesting that the ability to bind cholesterol was necessary but not sufficient for the cholesterol efflux function of NPC2 (Ko et al., 2003). Indeed, we later demonstrated that NPC2-mediated LE/LY cholesterol egress is likely due to its cholesterol transport activity, which involves direct contact between the surface of NPC2 and membranes (Cheruku et al., 2006; Xu et al., 2008), and that the residues identified by Ko et al (2003) as being unable to ‘rescue’ sterol accumulation in NPC2 patient cells, despite normal binding, were in fact markedly defective in cholesterol transport (McCauliff et al., 2015). Of particular note was that residues which, when mutated, abrogated sterol transfer between NPC2 and membranes, mapped not to a single domain but rather to several surface domains (McCauliff et al., 2015). This suggested, in turn, that NPC2 may be able to interact with more than one membrane simultaneously, and we and others have in fact shown that NPC2 promotes membrane-membrane interactions (Abdul-Hammed et al., 2010; McCauliff et al., 2011; McCauliff et al., 2015). Importantly, mutants with reduced sterol transfer rates were unable to clear accumulated cholesterol from NPC2 deficient fibroblasts and were also deficient in promoting membrane-membrane interactions (McCauliff et al., 2015). Notably, all of these impairments occur despite normal cholesterol binding, indicating that certain regions of the NPC2 surface may interact with membranes to effectively transfer cholesterol (McCauliff et al., 2015).

Earlier, using model membranes, we demonstrated a remarkable, order of magnitude stimulation in NPC2 cholesterol transfer rates by the incorporation of lysobisphosphatidic acid (LBPA), also known as bis-monoacylglycerol phosphate (Cheruku et al., 2006; Xu et al., 2008). LBPA is a structural isomer of phosphatidylglyerol with an atypical phospholipid stereoconfiguration. It is localized primarily to inner LE/LY membranes and is thought to be involved not only in the formation of these internal membranes and their architecture, but also in the sorting and efflux of LE/LY components, including cholesterol (Gruenberg, 2003; Hullin-Matsuda et al., 2007; Kobayashi et al., 1998; Kobayashi et al., 1999). Incubation of BHK cells and macrophages with an anti-LBPA monoclonal antibody resulted in cholesterol accumulation in the LE/LY compartment, resembling the NPC phenotype (Delton-Vandenbroucke et al., 2007; Kobayashi et al., 1999). Interestingly, Chevallier et al. showed that by enriching NPC1-deficient cells with exogenously added LBPA, the cholesterol accumulation was reversed (Chevallier et al., 2008). Based on the dramatic effects of LBPA on NPC2-mediated cholesterol transfer, however, we hypothesized a specific functional interaction between LBPA and NPC2, such that LBPA enrichment of cells deficient in NPC2 would not reverse cholesterol accumulation, in contrast to NPC1-deficient cells.

In the present studies, we demonstrate the direct interaction of NPC2 with LBPA, identify the likely LBPA-sensitive domain on the NPC2 surface, and establish the essential functional nature of NPC2-LBPA interactions in cholesterol egress from the LE/LY compartment. The results indicate the existence of a heretofore unknown NPC2-dependent, NPC1-independent pathway of cholesterol egress from the endosomal/lysosomal system that involves NPC2 interaction with LBPA. This in turn suggests that LBPA enrichment may be used to effect cholesterol egress in cells with defective NPC1 but intact NPC2.

## Results

### Predicted orientation of NPC2 in membranes

Our previous kinetics analyses strongly suggested that the mechanism of cholesterol transfer between NPC2 and membranes was via protein-membrane interaction (Cheruku et al., 2006; McCauliff et al., 2015; Xu et al., 2008). NPC2 does not contain any apparent transmembrane domains, nor are there experimentally documented membrane interactive domains to date. For *de novo* predictions we therefore employed the Orientation of Proteins in Membranes (OPM) Database, a curated online resource that predicts the spatial positions of known protein structures relative to the hydrophobic core of a lipid bilayer (Lomize et al., 2012). Using the crystal structure of bovine NPC2 (PDB ID: 1NEP), a loop domain consisting of hydrophobic residues as highlighted in **Figure 1**, was predicted to be highly membrane interactive, with a ΔG of −4.6 kcal/mol. This domain corresponds to 56-HGIVMGIPV-64 and consists primarily of the hydrophobic residues I58, V59, M60, I62, P63, and V64, plus the non-polar residues G61 and G57. Structurally, this domain forms a “hydrophobic knob” which presents prominently on the surface of the NPC2 protein. OPM predicts that this knob domain inserts into the hydrophobic space of the membrane model, positioning the sterol binding pocket of NPC2 near the membrane surface (**Figure 1A**) and, thus, in proximity to membrane sterols. These computational observations suggest that this knob domain may play a key role in the mechanism by which NPC2 is able to transport cholesterol between inner membranes of the LE/LY compartment. Of note, and as shown in **Figure 1B**, the primary sequence of the hydrophobic knob domain is conserved in mammalian NPC2 proteins but not in the yeast NPC2 homologue. Hydrophobicity scales for NPC2 as well as a Kyte-Doolittle plot of the hydrophobicity scores along the primary protein sequence indicate that the hydrophobicity of the knob domain is conserved amongst mammalian NPC2 proteins, in contrast to the low hydrophobicity of the yeast NPC2 protein (**Figure 1C**).

**Figure 1.**
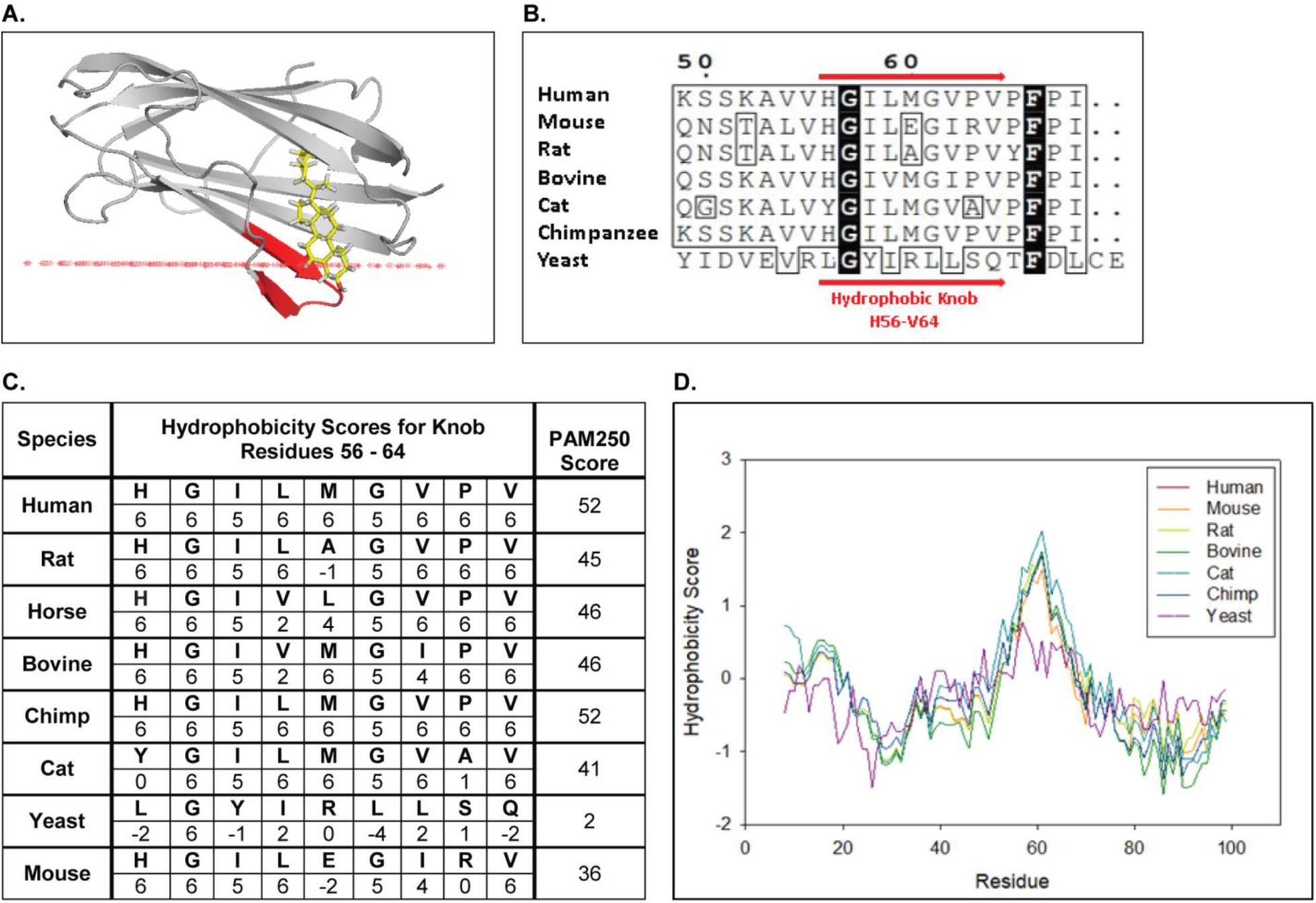
OPM predicts NPC2 stably inserts into membranes via a conserved hydrophobic knob region. (**A**) Holo bovine NPC2 (PDB ID: 2HKA) is predicted by OPM to insert its prominent hydrophobic knob region (red ribbon) into the hydrophobic space of a model membrane, positioning the cholesterol in the sterol binding pocket in close proximity to the membrane surface (red line). (**B**) Multiple sequence alignment of human NPC2 (NCBI Accession: NP_006423.1), rat NPC2 (NP_775141.2) mouse NPC2 (NCBI Accession: NP_075898.1), bovine NPC2 (NCBI Accession: NP_776343.1), cat NPC2 (XP_003987882.1), chimpanzee NPC2 (NP_001009075.1) and the yeast NPC2 (NCBI Accession: KZV12184.1) were aligned with CLUSTAL Omega, and alignment for hydrophobic knob residues H56 to V64 are shown. Consensus sequences are in black and conserved residues are boxed. (**C**) Conservation scores and hydrophobicity scores for the hydrophobic knob, residues H56 to V64, were calculated based on the PAM250 scoring matrix and Kyte & Doolitle Hydrophobicity scale (Kyte & Doolittle, 1982). (**D**) The aligned NPC2 protein sequences were analyzed with the ProtScale Tool on ExPASy server based on the Kyte and Doolittle Amino acid Hydropathicity scale with a frame window of 15 residues (Gasteiger et al., 2005).

### LBPA markedly stimulates sterol transfer rates between NPC2 and membranes

We previously found that incorporation of 25 mol% LBPA in EPC membranes resulted in cholesterol transfer rates from vesicles to NPC2 that were markedly accelerated relative to 100% EPC membranes (Cheruku et al., 2006; Xu et al., 2008). LBPA accounts for approximately 15 mol% of total LE/LY phospholipids, with the likelihood of higher lateral concentrations in the highly heterogeneous inner LE/LY membranes (Kobayashi et al., 2002). Thus, we examined the rates of cholesterol transfer from NPC2 to membranes as a function of increasing levels of LBPA. The results in **Figure 2** indicate an exponential relationship between the LBPA content of the vesicles and the NPC2 cholesterol transfer rate; increasing the mol% of LBPA in SUV from 0 to 30% effectively increases the NPC2 cholesterol transfer rates by approximately 100 fold.

**Figure 2.**
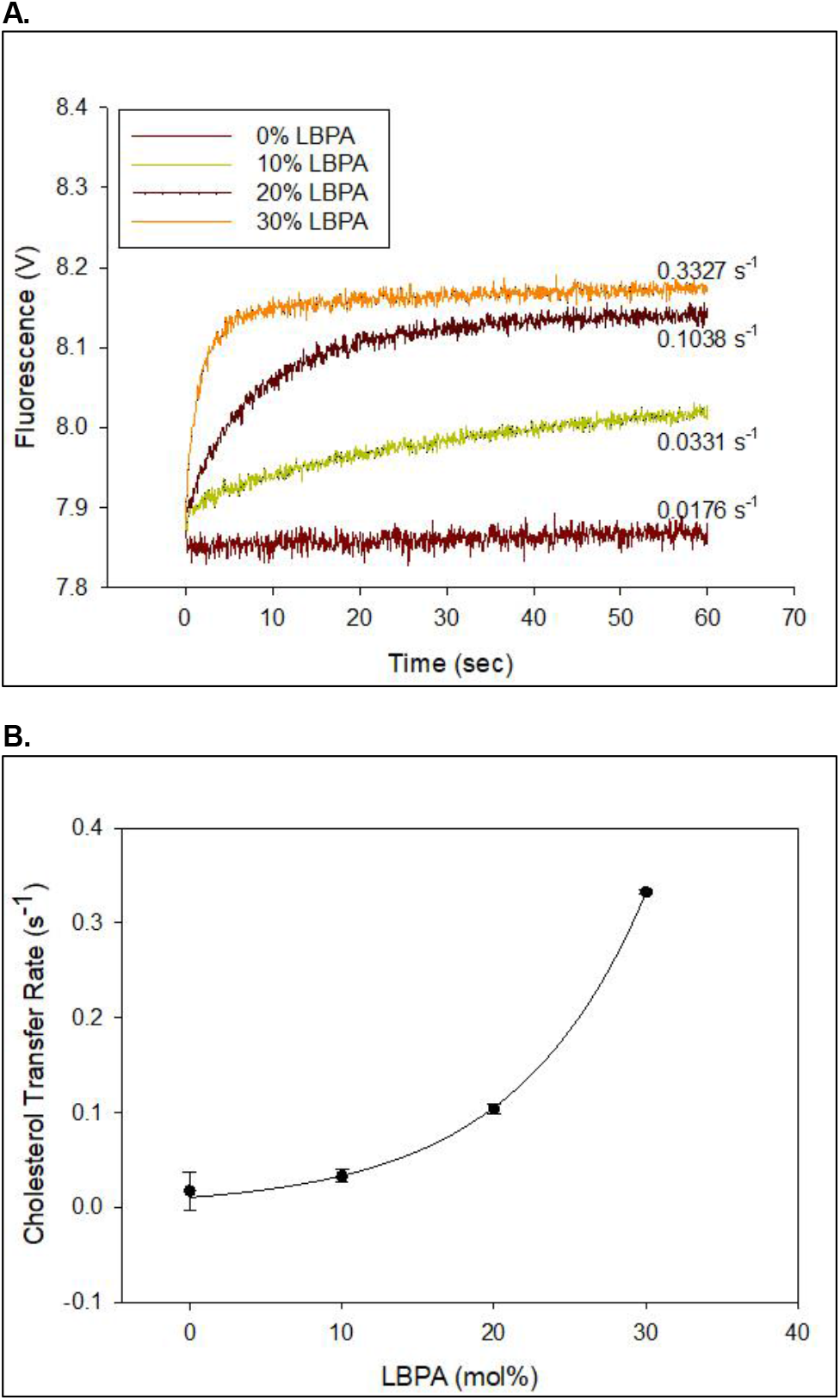
LBPA dramatically increase the rate of NPC2 mediated cholesterol transport. (**A**) 2.5 μM NPC2-cholesterol complex was mixed with 250 μM of small unilamellar vesicles containing increasing mole percentages of LBPA in an SX20 stopped flow spectrofluorimeter as described under Methods. The dequenching of endogenous tryptophan fluorescence was used to monitor cholesterol transfer from NPC2 to membranes. (**B**) The transfer rates from NPC2 to small unilamellar vesicles with various mole percentages of LBPA were fitted to a reverse single exponential function in Sigmaplot with an R-squared value of 0.9944. Data are representative of three experiments, each consisting of 2-3 individual runs ± SE.

### LBPA restores normal NPC2 cholesterol transfer rates for proteins with mutations outside the hydrophobic knob

We recently reported that point mutations in multiple regions on the NPC2 surface led to markedly diminished rates of cholesterol transfer between NPC2 and zwitterionic PC membranes; the hydrophobic knob domain was one of the regions found to be dramatically impacted (McCauliff et al., 2015). Here we examined the rates of cholesterol transfer from these and additional hydrophobic knob domain point mutations to membranes containing 25 mol% LBPA. Point mutations were confirmed by DNA sequencing and all mutant proteins were found to bind cholesterol similar to WT NPC2 (Friedland et al., 2003; Ko et al., 2003), with submicromolar affinity (McCauliff et al., 2015). The results in **Figure 3A** show that when acceptor membranes included 25 mol% of LBPA, cholesterol transfer rates for NPC2 proteins with mutations in regions *other* than the hydrophobic knob were similar to those of WT.

**Figure 3.**
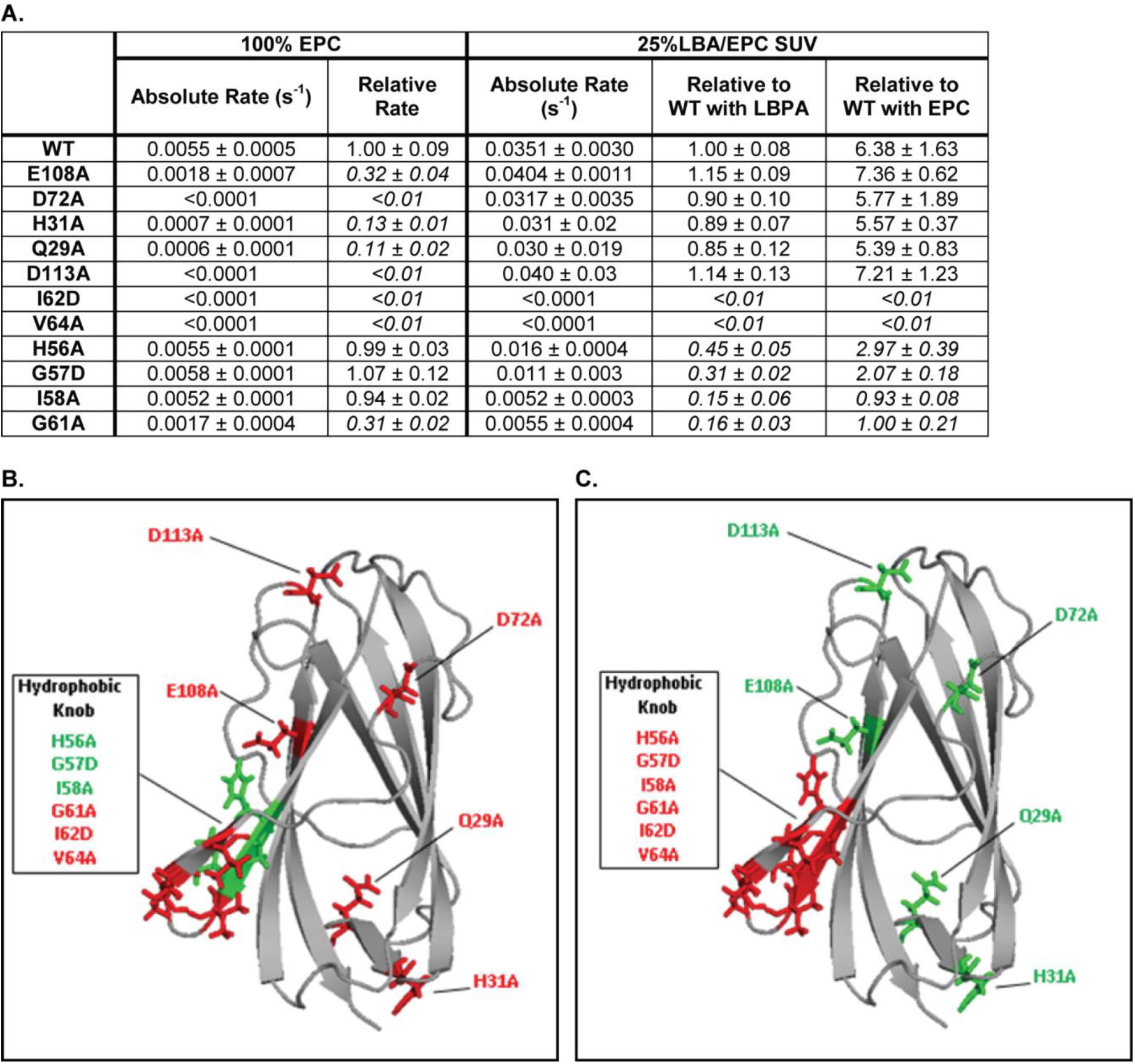
LBPA cannot reverse cholesterol transport deficiencies of NPC2 hydrophobic knob mutants. **(A)** Transfer of cholesterol from 1 μM WT or mutant NPC2 to 125 μM 100% EPC or 25% LBPA/EPC vesicles was measured on an SX20 Stopped Flow Spectrofluorometer by monitoring the dequenching of NPC2 endogenous tryptophan fluorescence. All curves were well fit using a single exponential function using the Applied Photophysics Pro-Data Viewer software. Mutants with rates of cholesterol transfer less than 50% of WT NPC2 were considered to have defective transfer kinetics properties and their relative rates are italicized. Data are representative of 3 experiments, each consisting of 2-3 individual runs. Absolute and relative rates of transfer to each model membrane system, ±SE, are shown. **(B)** NPC2 point mutations resulting in defective cholesterol transport to 100% EPC vesicles are shown in red while mutations having little or no effect on NPC2 cholesterol transport properties relative to WT protein, are shown in green. **(C)** Point mutations with attenuated rates of cholesterol transport to 25% LBPA/EPC vesicles are shown in red while mutations having little or no effect, relative to WT protein, are shown in green.

Indeed, though mutations at H31, Q29, D113, and E108 exhibited sterol transfer rates to EPC membranes that were ≤ 15% of WT NPC2, the inclusion of LBPA in acceptor membranes resulted in rates of cholesterol transfer that were ≥ 85% of WT rates. By contrast, the I62 and V64 mutations in the hydrophobic knob, which also resulted in markedly defective cholesterol transfer to EPC vesicles, were unaffected by the inclusion of LBPA in the acceptor membranes, with cholesterol transfer by these mutants remaining barely detectable. The G61A mutation, also in the hydrophobic knob, resulted in cholesterol transfer deficiencies similar to the I62 and V64 mutants, though changes are less extreme; sterol transfer to EPC vesicles was reduced by 70%, and it remained highly defective in the presence of LBPA, with rates of cholesterol transfer of only 16% relative to WT NPC2. Mutations in hydrophobic knob residues H56, G57, and I58 had little effect on cholesterol transfer rates to EPC vesicles, however, unlike WT NPC2, these mutants were insensitive to the presence of LBPA in acceptor membranes. The NPC2 residues where mutations cause large decreases in cholesterol transfer rates to EPC, are shown in red in **Figure 3B**; multiple surface regions are highlighted. By contrast, **Figure 3C** shows the mutations that were sensitive to LBPA inclusion, in green, vs. mutations which remained insensitive to membrane LBPA, in red; the hydrophobic knob is clearly denoted as LBPA-sensitive. Overall, the mutagenesis results strongly indicate that the hydrophobic knob domain of NPC2 is the LBPA-sensitive region on the protein surface.

### Effects of NPC2 mutations on vesicle-vesicle interaction also highlight the hydrophobic knob domain

We and others have shown that WT NPC2 promotes membrane-membrane interactions (Abdul-Hammed et al., 2010; Berzina et al., 2018; McCauliff et al., 2011; McCauliff et al., 2015). We further showed that NPC2 point mutants with deficient cholesterol transfer abilities are also unable to cause EPC membrane aggregation (McCauliff et al., 2015). In the present studies, we investigated whether the presence of LBPA in the vesicles affected vesicle aggregation by WT and mutant NPC2 proteins. The results in **Table 1** show that inclusion of 25 mol% LBPA in LUVs resulted in a 16-fold increase in the rate of membrane-membrane interaction by WT NPC2, relative to 100% EPC LUVs. Incorporation of LBPA into membranes normalizes the membrane aggregation rates for NPC2 proteins with mutations outside the hydrophobic knob, e.g. H31, D113, and Q29. By contrast, the hydrophobic knob domain mutations were relatively insensitive to membrane LBPA. The results for this membrane-membrane interaction assay map virtually identically onto the NPC2 structure as did those for the cholesterol transport rates, as seen in **Figure 3**, again indicating that the hydrophobic knob of NPC2 is the LBPA-sensitive region of the protein.

**Table 1.** Rate of NPC2-mediated membrane interaction greatly increases in the presence of LBPA. The effect of surface residue mutations on the ability of NPC2 to induce vesicle-vesicle interactions was assessed by measuring absorbance at 350nm (light scattering) of 200 μM LUVs in the presence of 1 μM WT or mutant NPC2 protein, as described under Methods. Rates of vesicle-vesicle interactions, indicated by increases in A350nm over time, were determined by a three-parameter hyperbolic fit of the data using Sigma Plot software, and are representative of at least three individual experiments. Mutants with substantially attenuated rates of membrane aggregation are indicated in italics.

|  | 100% EPC |  | 25%LBA/EPC SUV |  |  |
| --- | --- | --- | --- | --- | --- |
|  | Absolute Rate (s<sup>-1</sup>) | Relative Rate | Absolute Rate (s<sup>-1</sup>) | Relative to WT with LBPA | Relative to WT with EPC |
| WT | 0.0112 ± 0.0006 | 1.00 ± 0.05 | 0.1674 ± 0.0122 | 1.00 ± 0.07 | 14.95 ± 1.09 |
| H31A | 0.0018 ± 0.0005 | *0.16 ± 0.05* | 0.1262 ± 0.0079 | 0.75 ± 0.05 | 11.27 ± 0.71 |
| D113A | <0.0001 | *<0.01* | 0.1573 ± 0.0112 | 0.94 ± 0.07 | 14.04 ± 1.00 |
| Q29A | 0.0029 ± 0.0008 | *0.26 ± 0.07* | 0.1337 ± 0.0039 | 0.80 ± 0.02 | 11.93 ± 0.35 |
| E108A | 0.0032 ± 0.0004 | *0.29 ± 0.03* | 0.1437 ± 0.0094 | 0.86 ± 0.06 | 12.83 ± 0.84 |
| D72A | 0.0032 ± 0.0003 | *0.29 ± 0.03* | 0.1456 ± 0.0077 | 0.87 ± 0.05 | 13.00 ± 0.69 |
| H56A | 0.0107 ± 0.0008 | 0.95 ± 0.07 | 0.0261 ± 0.0055 | *0.16 ± 0.09* | *2.33 ± 0.49* |
| G57D | 0.0103 ± 0.0016 | 0.92 ± 0.14 | 0.0202 ± 0.0066 | *0.13 ± 0.12* | *1.81 ± 0.59* |
| I58A | 0.0094 ± 0.0011 | 0.840 ± 0.10 | 0.0240 ± 0.0037 | *0.15 ± 0.06* | *2.14 ± 0.33* |
| G61A | 0.0075 ± 0.0006 | *0.67 ± 0.06* | 0.0406 ± 0.0012 | *0.25 ± 0.03* | *3.63 ± 0.11* |
| I62D | <0.0001 | *<0.01* | 0.0167 ± 0.0017 | *0.10 ± 0.03* | *1.50 ± 0.15* |
| V64A | <0.0001 | *<0.01* | 0.0193 ± 0.0040 | *0.12 ± 0.06* | *1.72 ± 0.36* |

### NPC2 interaction with LBPA and other phospholipids, and identification of the LBPA-interactive domain

To determine whether the relationship between LBPA and NPC2 in cholesterol trafficking involves direct interactions, protein-lipid binding assays were conducted. For studies of WT NPC2 protein, custom LBPA Snoopers (Avanti Polar Lipids) containing various LBPA isomers were incubated with WT NPC2 protein and relative binding was assessed via densitometric analysis of an antibody-probed strip, as described in Methods. The results shown in **Figure 4A** demonstrate that WT NPC2 binds to LBPA, showing greater interaction with isomers containing oleoyl (C18:1) as opposed to myristoyl (C14:0) fatty acyl chains. Interestingly, WT NPC2 exhibited the greatest degree of binding to the *S,S* 18:1 LBPA, which possess not only the likely stereoconfiguration in mammals (Chevallier et al., 2000; Hayakawa et al., 2006; Kobayashi et al., 2002), but also the predominant fatty acyl chains in most cell types studied thus far (Kobayashi et al., 2002; Mason et al., 1972). NPC2 binding of the *S,S* 18:1 LBPA was also greater than binding to egg PC.

Binding of WT NPC2 to other typical membrane phospholipid species, in comparison to di-oleoyl LBPA, was also examined using membranes spotted with di-oleoyl phospholipids. **Figure 4B** shows that NPC2 interacts more strongly with LBPA than with PC, PA, PG, and PS species. As two other anionic phospholipids assayed, PG and PA, show even less interaction with NPC2 than does zwitterionic PC, the mechanism by which NPC2 binds to LBPA is likely not solely dependent on electrostatic interactions.

**Figure 4.**
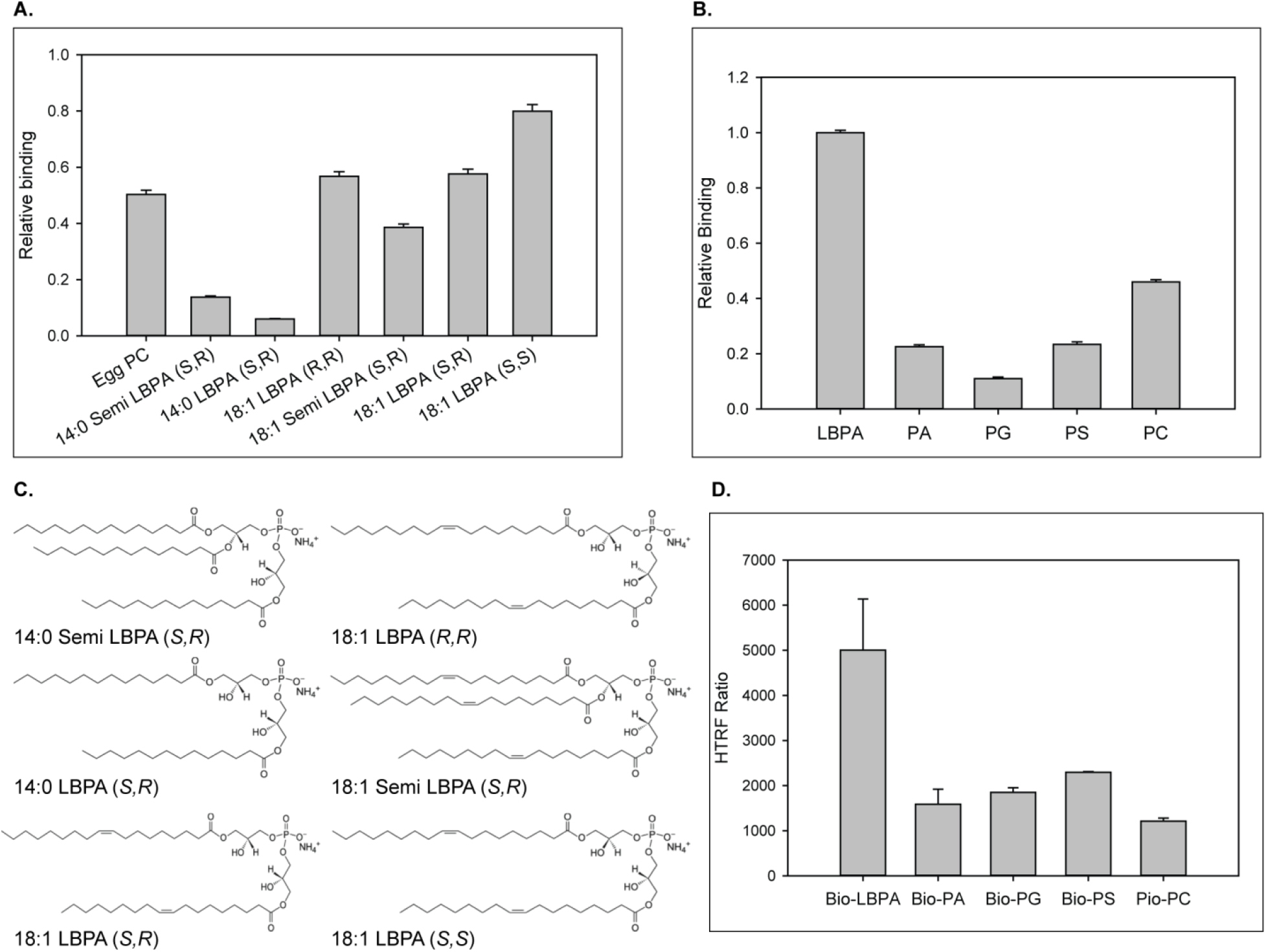
NPC2 binding to LBPA and other phospholipids. (**A**) WT NPC2 protein was incubated with strips (Snoopers) containing LBPA isomers. LBPA-bound protein was detected with anti-c-myc antibody as described under Methods, and degree of binding ± SE (n=3) is represented by the integrated density of the blots. (**B**) 500 pmol of various membrane phospholipids were spotted onto nitrocellulose strips and probed with WT NPC2-myc-his protein as described under Methods. Relative binding of WT NPC2 ± SE (n=5) is shown, represented by signal intensity detected with the LI-COR system. (**C**) Structures of the LBPA isomers. (**D**) 75nM of WT NPC2 protein was incubated with 1μM of the indicated biotin-C12-ether phospholipid, streptavidin-d2 conjugate and europium cryptate–labeled monoclonal anti-histidine antibody in detection buffer, as described in Methods. FRET signal between europium cryptate and streptavidin was detected with a HTRF capable Envision plate reader (λex = 320 nm, λem = 615 and 665 nm; 100 μs delay time; n=3).

Since the phospholipids are not necessarily present in a physiological orientation in protein-lipid overlay assays, we further examined NPC2–lipid interaction using Homogenous Time Resolved Fluorescence (HTRF), in which the phospholipids are present as lamellar structures. HTRF technology has been recently demonstrated to be an effective and sensitive assay for lipid-protein interaction (Fleury et al., 2015). The NPC2-interaction with LBPA was substantially greater than with other negatively charged lipids or with zwitterionic PC (**Figure 4D**), in general agreement with the lipid-blot results. Taken together the results support direct interactions between NPC2 and membrane phospholipids, the greatest interaction being with LBPA.

To determine whether there is a specific LBPA interactive domain on the surface of NPC2, we further employed the LBPA lipid blots to examine interactions with NPC2 point mutants. The results show that mutations in most of the hydrophobic knob residues markedly reduce interactions, relative to WT NPC2. For example, studies analyzing binding to various isomers of LBPA show that the hydrophobic knob mutants I62N and V64A bound the *S,S* and *S,R* di-oleoyl isomers at only ~ 20% of WT levels, and the C18:1 *R,R* isomer at approximately 40% of WT. In contrast, the Q29A and D113A mutants, in regions outside the hydrophobic knob, bound nearly all LBPA isomers similar to WT; only Q29A was observed to bind the C18:1 *R,R* isomer at approximately 50% relative to WT binding (**Figure 5A**). The I62D and V64A mutants exhibited greater interaction with the 18:1 Semi LBPA species, which has three oleoyl acyl chains (**Figure 5A**), binding approximately 70% relative WT NPC2. Mutations in other hydrophobic knob residues also reduced the interaction of NPC2 with the di-oleyol *S,S* LBPA on blots using various phospholipids. Indeed, similar to what was observed with the LBPA isomer blots, the H56, G57, I58, and G61 mutants showed only 30 to 40% degree of interaction relative WT NPC2. In marked contrast, surface mutations outside the hydrophobic knob had virtually no impact on NPC2 interaction with LBPA; Q2A, H31A, D113, and E108 mutants exhibiting WT levels of interaction with LBPA (**Figure 5B**). Overall, mutations within the hydrophobic knob domain of NPC2 (H56, G57, I58, G61, I62 and V64) resulted in diminished binding of the protein to LBPA while mutations outside the knob region (Q29, H31, E108, D113) presented proteins with LBPA interactions similar to WT (**Figure 5B**). HTRF analysis of the mutant NPC2 proteins was generally consistent with the lipid blot results, also indicating reduced binding of the hydrophobic knob mutants to LBPA (**Figure 5C,D**). Thus, the results identify the hydrophobic knob of NPC2 as the LBPA-interactive domain on the protein.

**Figure 5.**
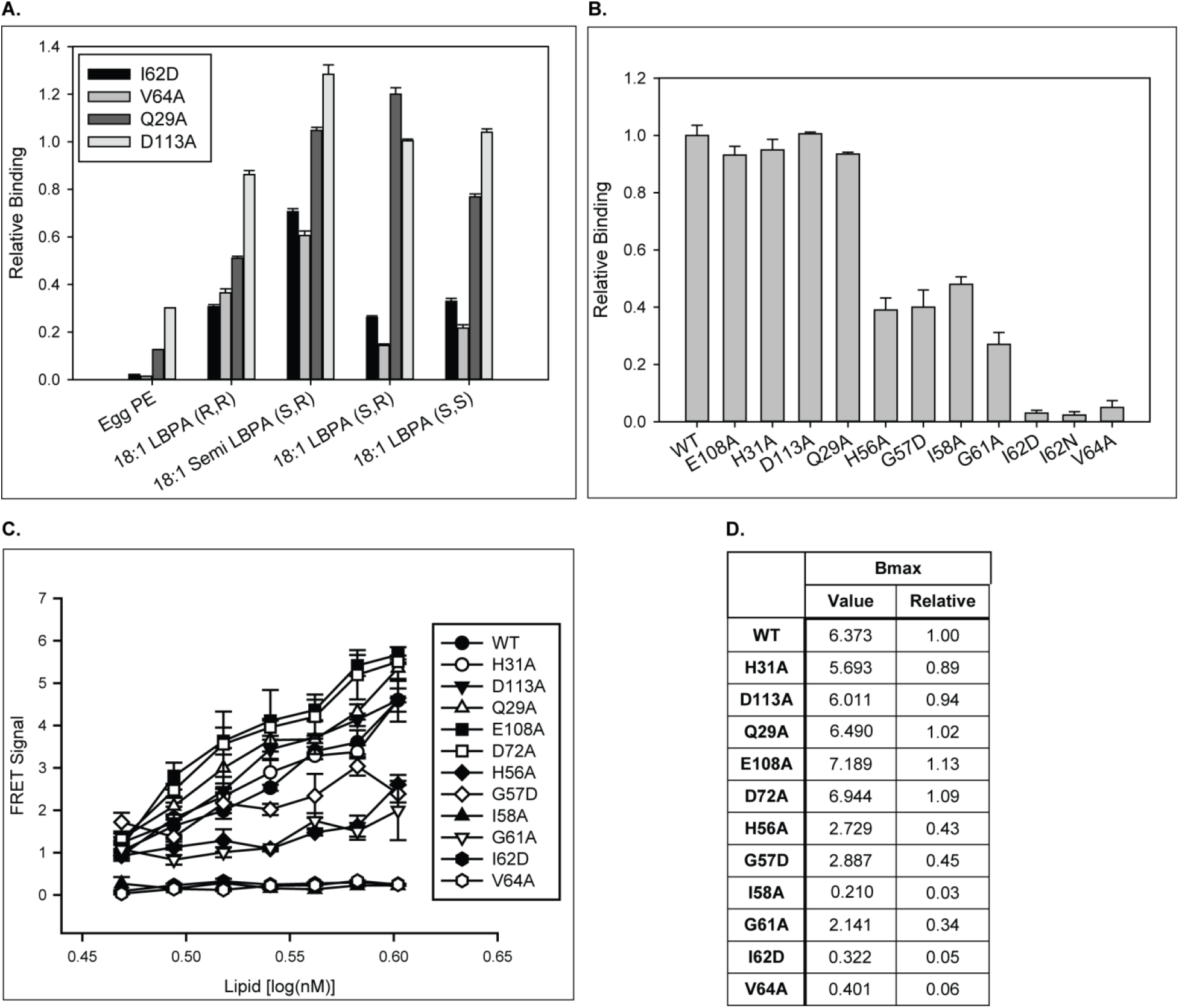
NPC2 binds to LBPA via the “hydrophobic-knob” domain. Binding of NPC2 WT and mutant proteins to (**A**) LBPA isomers and (**B**),18:1 (S,R) LBPA was detected using LBPA Snoopers and membranes spotted with 500 pmol phospholipid, respectively, as described under Methods. Relative binding is represented as (A) the integrated density of the blots, relative to WT NPC2, ±SE (n=3) and (B) the signal intensity detected with the LI-COR system ±SE (n=3) (**C**) HTRF analysis: WT or mutant NPC2 protein was incubated with biotin-C12-ether LBPA, streptavidin-d2 conjugate and europium cryptate-labeled anti-His antibody in detection buffer, as described in Methods. FRET signal between europium cryptate and streptavidin was detected with a HTRF capable Envision late reader as described in Methods. (**D**) FRET signal was analyzed with the One site–specific binding function (Graphpad) and the Bmax extrapolations were used to infer binding capacity between recombinant NPC2 protein and biotin-C12-ether LBPA. Results are representative of three experiments, with deviations ≤20%.

### Transfer kinetics and LBPA interaction predict ability of WT or mutant NPC2 to reduce cholesterol accumulation in NPC2 deficient fibroblasts

In agreement with several previous reports, incubation of NPC2 deficient fibroblasts with WT NPC2 protein resulted in a dramatic decrease in filipin staining, reaching levels similar to healthy fibroblasts (**Figure 6**) (Ko et al., 2003; Liou et al., 2006; McCauliff et al., 2011; McCauliff et al., 2015). The H56A mutant, with rates of sterol transfer and membrane aggregation similar to WT protein, also reduced filipin stain area, similar to WT NPC2. In contrast, the G57D and I58A hydrophobic knob mutants, with markedly attenuated cholesterol transfer and membrane aggregation rates, were unable to reverse cholesterol accumulation in NPC2 cells; filipin staining remained at a level comparable to that of the unsupplemented cells. G61A, also in the knob domain, was able to lessen cholesterol accumulation in NPC2 cells to a moderate extent, and its defect in cholesterol transfer was also more modest than that of other hydrophobic knob mutants (**Figure 6**). These results are in keeping with our previously reported results for two other hydrophobic knob mutants, I62D and V64A (McCauliff et al., 2015) which are also deficient in cholesterol transfer and membrane aggregation ability. The consistency between results of the sterol transfer assays and membrane-membrane interaction assays, with the cholesterol clearance in patient cells, strongly supports the physiological relevance of the structure-function studies, and points to a particularly important role for the hydrophobic knob of NPC2 in effecting normal sterol trafficking.

**Figure 6.**
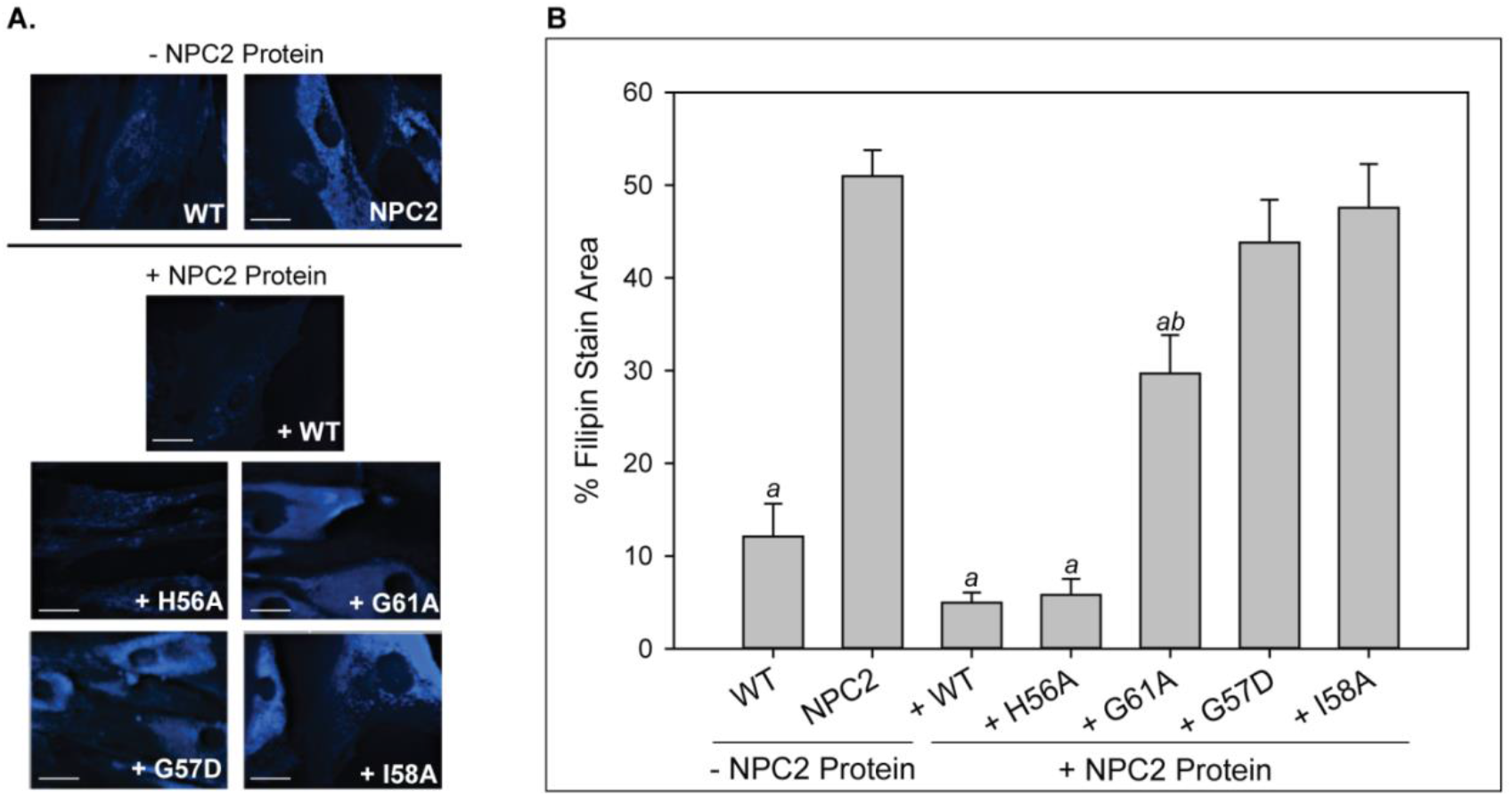
LBPA insensitive mutants are unable to rescue cholesterol accumulation in patient fibroblasts. NPC2-deficient fibroblasts were incubated with 0.4nm purified WT or NPC2 mutant protein and cholesterol accumulation was quantified via filipin staining as described in the Methods. (**A**) Representative microscopy images of filipin stained control and treated NPC2-deficient fibroblasts. Scale bars, 70μM. (**B**) Percent filipin stain area of untreated control and treated NPC2-deficient fibroblasts was quantified. Data are representative of at least three separate experiments ± SE. a, p<0.01 vs untreated NPC2-deficient cells; b, p<0.01 vs untreated WT cells by Student’s *t*-test.

### PG supplementation Increases LBPA levels and reduces cholesterol accumulation in NPC1-but not NPC2-deficient fibroblasts

In 2008, Chevallier et al. showed that supplementation of NPC1 deficient cells with LBPA via viral mediated delivery resulted in a marked reduction of intracellular cholesterol levels (Chevallier et al., 2008). However, based on the dramatic effects of LBPA on NPC2 cholesterol transfer and membrane-membrane interaction rates, and identification of the LBPA interaction domain on the NPC2 surface, we predicted that LBPA enrichment of NPC2 deficient cells would *not* ameliorate the cholesterol accumulation phenotype.

To increase cell LBPA levels, NPC patient fibroblasts were incubated with PG, known to be its precursor (Bouvier et al., 2009; Poorthuis & Hostetler, 1978; Thornburg et al., 1991). There is currently some uncertainty as to the stereoconfiguration and acyl chain attachment site of LBPA in cells (Chevallier et al., 2008; Goursot et al., 2010; Hayakawa et al., 2006), thus we reasoned that the endogenous synthesis of LBPA from its precursor PG would allow for generation of the appropriate LBPA isomer, providing a physiologically relevant model. The results in **Figure 7** show that incubation of the cells with 100% PG SUVs led to substantial increases in cellular content of LBPA in all fibroblast types; a nearly 6-fold increase was observed in WT cells. In agreement with previous reports LBPA levels in NPC patient cells were found to be increased relative to WT cells prior to enrichment (Chevallier et al., 2008; Davidson et al., 2009; Sleat et al., 2004; Vanier, 1983); PG incubation resulted in 2-3 fold increases in NPC1- and NPC2-deficient cells, respectively, relative to unsupplemented cells (**Figure 7A**). Interestingly, the levels of other cellular phospholipids remained unchanged, including PG itself (**Figure 7B**). Increases in LBPA were also visualized using the 6C4 anti-LBPA antibody, as described in Methods, and results showed 2-fold or greater increases in LBPA levels relative to unsupplemented cells (**Figure 7C**). Direct addition of LBPA SUVs also led to approximately 2 to 3-fold increases in cellular LBPA levels (data not shown), in agreement with Chevallier et al (Chevallier et al., 2008). Supplementation with PC SUVs as a control had no effect on the phospholipid composition of any of the cell types (data not shown).

**Figure 7.**
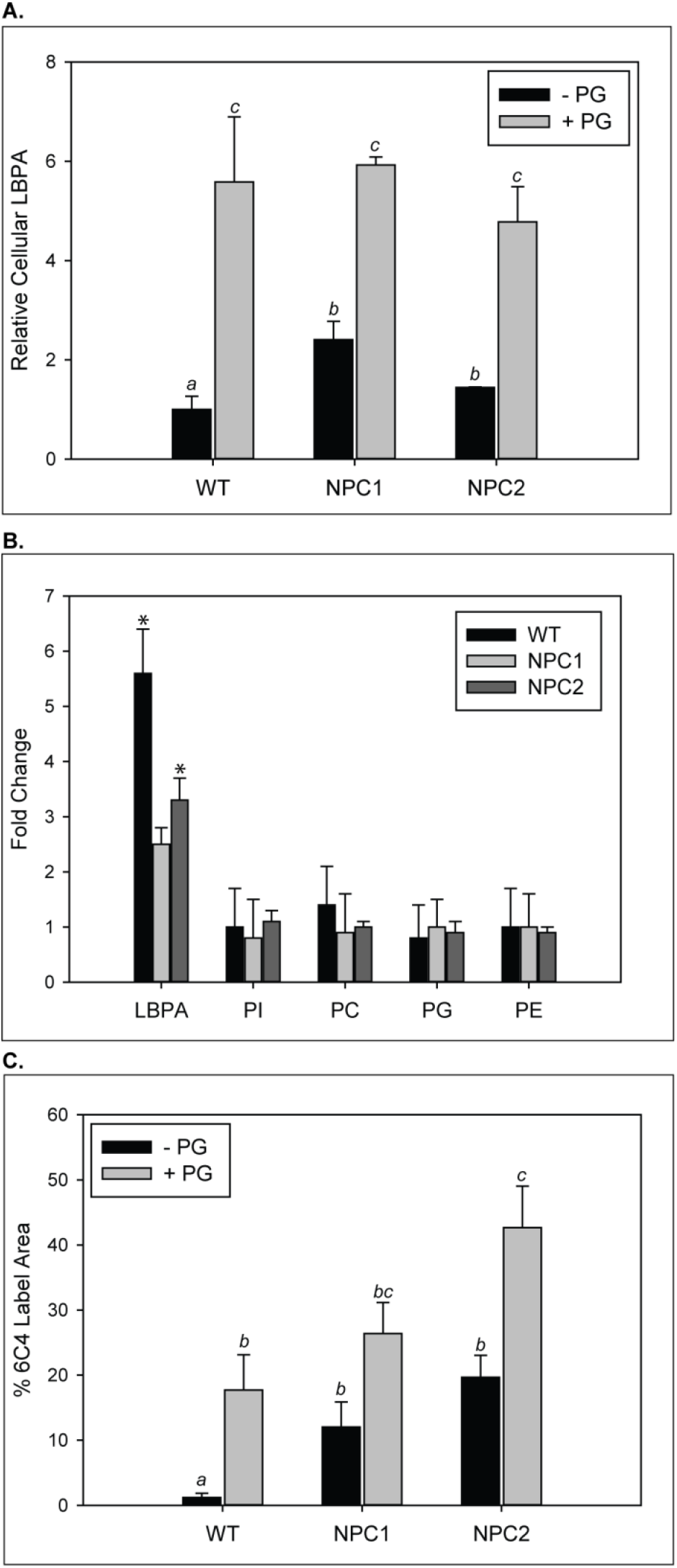
Increase in LBPA content in cells supplemented with PG. WT, NPC1-, and NPC2 deficient fibroblasts were incubated with 100 μM PG SUVs. (**A-B**) Lipids were extracted and phospholipids quantified by TLC as described under Methods. Results are representative of two experiments, each conducted in duplicate, ± SE. (**A**) Data are normalized to LBPA levels in untreated WT cells. (**B**) Fold changes in PL species in PG-treated relative to untreated cells; *p<0.01 between PL species by one-way ANOVA. TLC results were confirmed by (**C**) quantification of 6C4 anti-LBPA antibody staining, as described under Methods. Data are representative of three individual experiments ± SE. a, p<0.01 vs untreated WT control cells; b, p<0.01 vs untreated NPC1-deficient cells; c, p<0.01 vs untreated NPC2-deficient cells by Student’s *t*-test.

We evaluated the effect of LBPA enrichment via PG supplementation on intracellular cholesterol content in WT and NPC disease fibroblasts using filipin staining. Following PG supplementation, cholesterol content remained apparently unchanged in the WT fibroblasts. NPC1 deficient fibroblasts exhibited a dramatic reduction in cholesterol accumulation following PG supplementation/LBPA enrichment, approaching levels observed for WT cells, similar to the direct addition of LBPA (Chevallier et al., 2008). In marked contrast, and as we hypothesized based on NPC2-LBPA interactions and the effects of LBPA on NPC2 sterol transfer rates, the cholesterol accumulation in NPC2 deficient fibroblast persisted following PG supplementation despite increased LBPA content (**Figure 8**). Thus, the results support the presence of an NPC2-LBPA dependent, NPC1-independent, mode of LE/LY cholesterol efflux.

**Figure 8.**
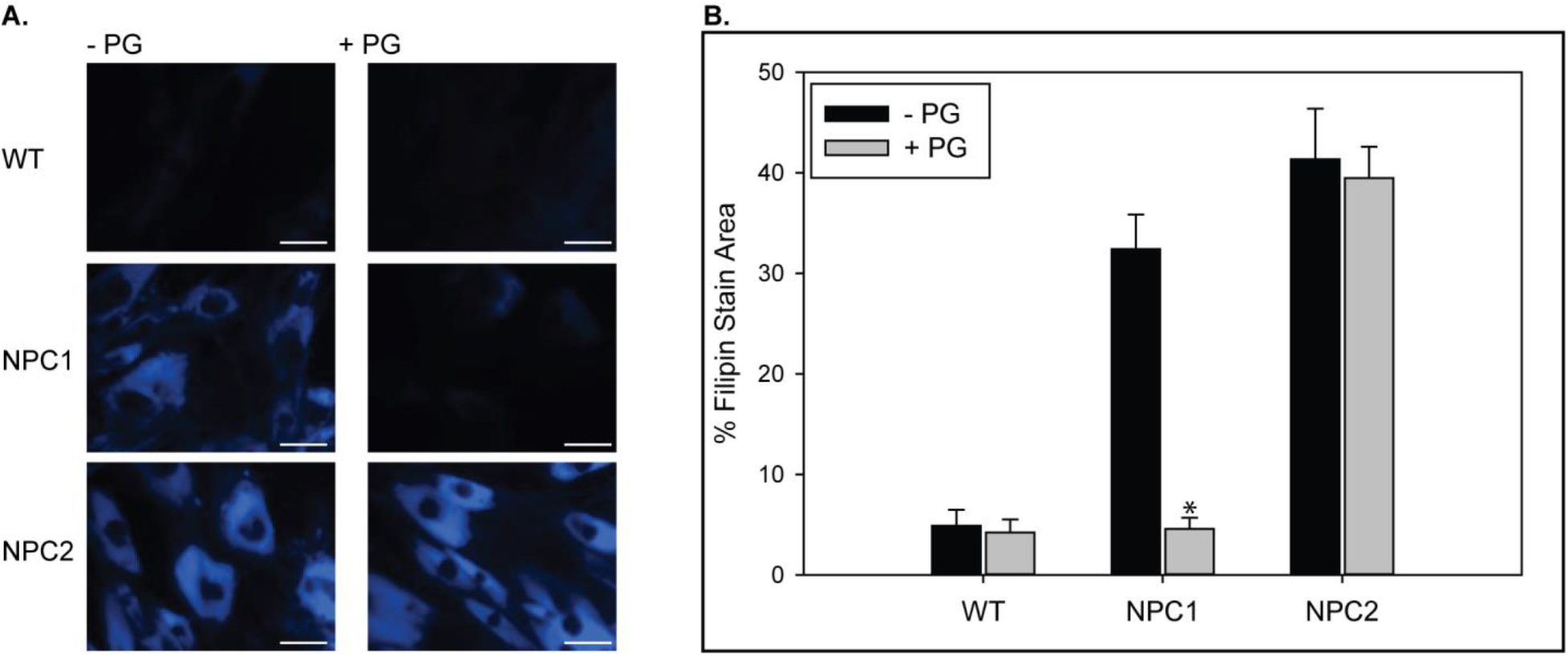
PG supplementation reverses cholesterol accumulation in NPC1– but not NPC2–deficient cells. WT, NPC1–, and NPC2–deficient fibroblasts were incubated with 100 μM PG SUVs and cholesterol accumulation was determined by filipin staining as described under Methods. **(A)** Representative images of untreated and PG supplemented fibroblasts stained with filipin. Scale bars, 70μM. **(B)** Percent of cell area stained with filipin. Data are representative of three individual experiments, ± SE. *p<0.01 vs untreated cells by Student’s *t*-test.

### Cellular LBPA accumulation restores cholesterol clearance properties of LBPA-sensitive NPC2 mutants

The point mutagenesis analysis of NPC2 shows a direct relationship between the cholesterol transfer rate of a particular NPC2 mutant and its ability to rescue the cholesterol accumulation of NPC2-deficient cells (McCauliff et al., 2015); **Figure 3 & 6**). In the present studies we discovered an unanticipated impact of LBPA incorporation into membranes, in which some NPC2 mutants that were highly defective in sterol transfer to phosphatidylcholine membranes, were essentially normalized when LBPA was present. These “LBPA-sensitive” NPC2 mutations were, almost exclusively, present in surface domains outside of the hydrophobic knob. NPC2 proteins with mutations in the hydrophobic knob, that were highly defective in cholesterol transfer to phosphatidylcholine membranes, remained insensitive to LBPA in the membranes (**Figure 3**). Based on this structure-based difference in LBPA sensitivity, we hypothesized that increasing the LBPA levels in NPC2 deficient cells would enhance the activity of mutants outside the hydrobphobic knob that responded to LBPA in the kinetic assays, whereas cellular LBPA enrichment would not enhance the action of the hydrophobic knob mutants that were insensitive to LBPA. To test these predictions, NPC2 deficient fibroblasts were incubated with purified wild type or mutant NPC2 proteins, with or without PG supplementation, and cholesterol accumulation was assessed by filipin staining as described. The results in **Figure 9** demonstrate that NPC2 proteins with mutations outside the hydrophobic knob, such as Q29A, D113A, and D72A, which were unable to reduce cholesterol accumulation in unsupplemented NPC2 deficient cells, were indeed able to ‘rescue’ cells that were enriched with LBPA via PG supplementation. By contrast, the hydrophobic knob mutants I62D and G61A, which were insensitive to LBPA in cholesterol transfer assays, were unable to clear cholesterol from LBPA-enriched NPC2 deficient cells. Thus, the results show that a combination of increased cellular levels of LBPA and LBPA-sensitive NPC2 protein was able to rescue the NPC2 deficient cells, beyond the ability of the mutant protein alone (**Figure 9**). As before, WT cells supplemented with PG alone showed no change in cholesterol accumulation, and cells supplemented with purified WT NPC2, with or without PG, showed significantly reduced filipin staining. The results in patient cells again mirror the results of sterol kinetics experiments, suggesting that return to normal sterol trafficking can be achieved for NPC2 mutations outside the hydrophobic knob that are otherwise dysfunctional, when the LBPA content of the cells is increased.

**Figure 9.**
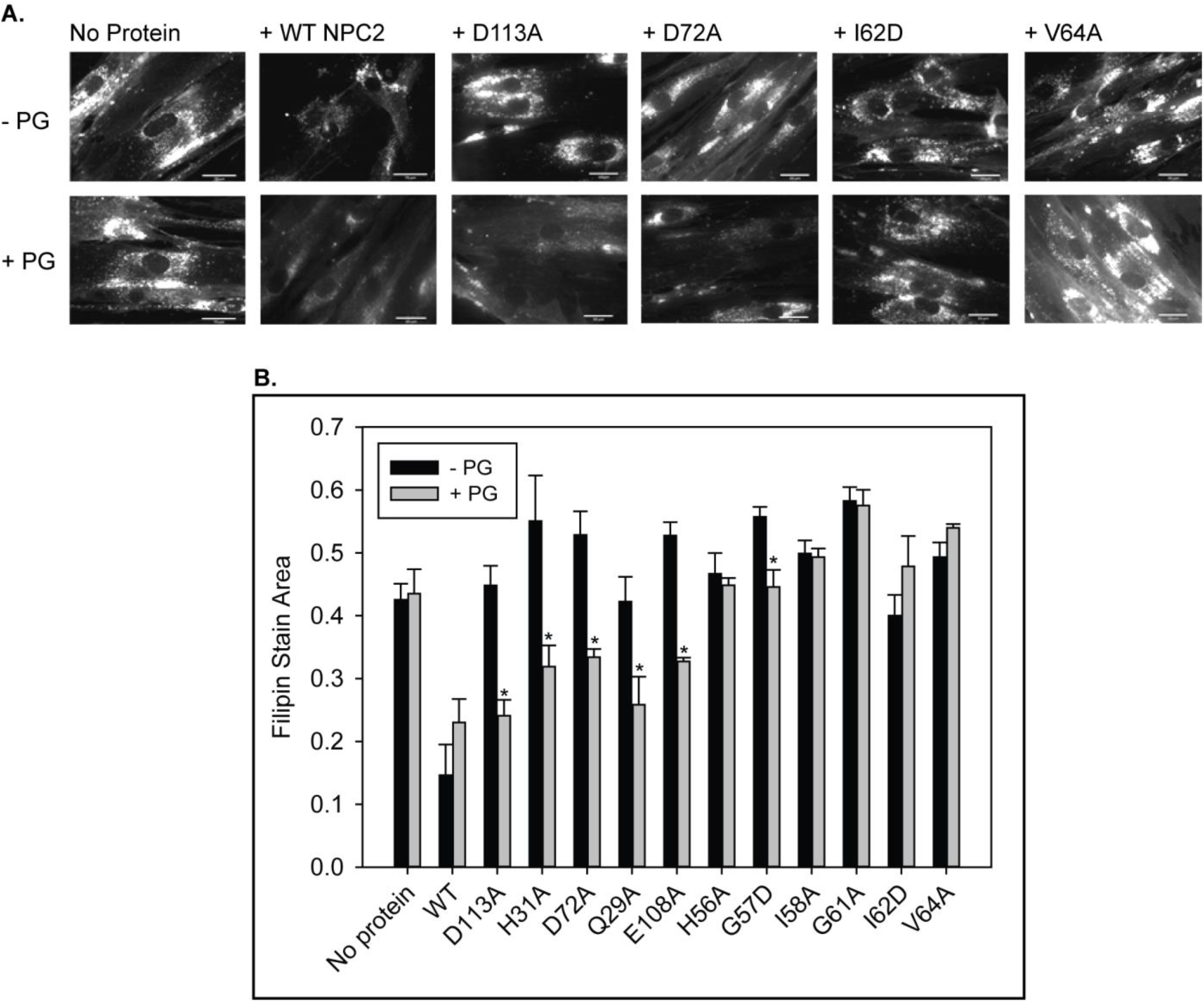
LBPA–sensitive but not insensitive NPC2 mutants reverse cholesterol accumulation in NPC2–deficient cells when co-treated with PG. NPC2 deficient fibroblasts were incubated with purified WT or mutant NPC2 proteins alone or in the presence of PG SUVs, as described under Methods, and cholesterol accumulation was quantified by filipin staining. (**A**) Representative images of treated NPC2 deficient fibroblasts stained with filipin. Scale bars, 70μM. (**B**) Data are from 4 or more individual incubations ± SE, *p<0.01 vs no PG supplementation by Student’s *t*-test.

## Discussion

LBPA was proposed to be involved in intracellular cholesterol trafficking based on the sterol accumulation which accompanies incubation of cells with an anti-LBPA antibody (Kobayashi et al., 1999), however the mechanism of LBPA action has remained unknown. In this study we demonstrate for the first time that functional and likely direct interaction of NPC2 protein with LBPA is a required step in normal cholesterol trafficking through the endosomal/lysosomal compartment. We show that NPC2 interacts with membrane LBPA, and that the hydrophobic knob domain is the site of NPC2-LBPA interaction. This surface domain is located near the NPC2 cholesterol binding pocket, thus its insertion into the bilayer would position the protein to efficiently exchange cholesterol with the membrane. Recent molecular dynamic simulations show that LBPA, but not other phospholipids, may position NPC2 in an orientation that could promote protein-membrane sterol exchange (Enkavi et al., 2017).

A decade ago, the Gruenberg laboratory reported that viral-mediated supplementation of NPC1-deficient cells with exogenous LBPA reversed cholesterol accumulation in the diseased cells (Chevallier et al., 2008). Here we show that LBPA enrichment of NPC2 deficient cells was completely ineffective, underscoring the required functional interaction of LBPA with the NPC2 protein. Nevertheless, however, we found that several surface mutations on NPC2 that could not reverse cholesterol accumulation in unsupplemented NPC2-deficient cells, became effective if the cells were first enriched with LBPA. The NPC2 mutations which are ‘rescued’ by LBPA enrichment are all localized in regions outside of the hydrophobic knob domain. By contrast, mutations within the hydrophobic knob were insensitive to LBPA enrichment with the exception of H56A, which may be on the limiting edge of this region. Thus, LBPA is able to normalize deficient transfer by mutant NPC2 as long as the mutation is outside of the hydrophobic knob. Interestingly, yeast lacks the LBPA phospholipid species. Primary amino acid sequence comparisons reveal that the yeast NPC2 exhibits a low degree of conservation within the hydrobhobic knob domain, and a markedly lower degree of hydrophobicity compared to all other species examined, supporting the notion that the hydrophobic knob domain is indeed critical to the protein’s functional relationship with LBPA.

LBPA is reported to comprise only 1% of total cellular phospholipids, but about 15 mol% of total phospholipids in the LE/LY (Chevallier et al., 2000; Kobayashi et al., 1998; Kobayashi et al., 2002). There is no agreement regarding which isomeric form of LBPA is predominant in cells, as the fatty acids can be linked at the sn-2 or sn-3 positions of each glycerol in the *S* or *R* conformation, though the sn-2, sn-2’ positions are currently favored (Chevallier et al., 2000; Kobayashi et al., 1998; Kobayashi et al., 2002; Mason et al., 1972; Matsuo et al., 2004). The 2,2’-LBPA was shown to be quite effective at mobilizing cholesterol in NPC1 disease cells while the 3,3’- and semi-LBPA isoforms were unable to promote sterol efflux. No difference in efficacy between *S,S, S,R*, and *R,R* LBPA isomers were noted, however, suggesting that the conformation of LBPA has no bearing on its ability to reverse cholesterol accumulation in NPC1 deficient cells (Chevallier et al., 2008). In agreement with this observation, we previously showed that the *S,S, S,R*, and *R,R* configurations of LBPA had little to no effect on *in vitro* cholesterol transfer rates by NPC2 protein (Xu et al., 2008). Similarly, in the present studies we observed little variation in NPC2 binding to these different LBPA stereoisomers. The cellular fatty acyl chain components of LBPA have also been found to vary (Bouvier et al., 2009), with oleic acid and docosahexaenoic acid (DHA) reported to be selectively incorporated (Besson et al., 2006; Luquain et al., 2001). Here we found acyl chain-dependent differences in NPC2-LBPA interactions, with reduced binding to the 14-carbon saturated dimyristoyl-LBPA species relative to the 18-carbon monounsaturated dioleoyl-LBPA species. In prior work we showed that cholesterol transfer from NPC2 to membranes containing dioleoyl-LBPA was > 2-fold faster than transfer to vesicles with dimyristoyl-LBPA (Xu et al., 2008). Taken together, the results suggest that the acyl composition but not the steroconfiguration of LBPA may be important in normal LE/LY cholesterol efflux.

In healthy cells, the enrichment of inner LE/LY membranes with LBPA and concurrent decline in cholesterol content may be reflective of the efficient role LBPA plays in normal LE/LY cholesterol efflux. If direct interactions between LBPA and NPC2 are integral to this mechanism of efflux, a rate determining step may be their frequency of interaction. While the concentration of LE/LY LBPA has been shown to increase in parallel with cholesterol and other lipids in NPC disease (Davidson et al., 2009; Sleat et al., 2004; Vanier, 1983), as also found here, it is possible that LBPA levels nevertheless remain too low to provide support for NPC2-mediated transport of the elevated cholesterol load. Gruenberg and colleagues noted the possibility of this limitation, showing that although cholesterol laden NPC1 cells had elevated levels of LBPA, virus-mediated supplementation of exogenous LBPA was effective at reversing the sterol accumulation phenotype (Chevallier et al., 2008). Our results using PG supplementation to augment LBPA levels further indicate that LBPA may become limiting in NPC1 disease. Importantly, we also show here that LBPA enrichment cannot reverse cholesterol accumulation caused by NPC2 deficiency.

In addition to the apparently obligate interaction of NPC2 with LBPA, LBPA has been found to display a variety of other unique functions within the endo/lysosomal system that could potentially promote this cooperative mechanism of cholesterol efflux. It has been shown to modulate phospholipid membrane curvature, for instance, and is required for formation of the multivesicular structures within the LE/LY (Matsuo et al., 2004), where cholesterol localizes in raft domains (Fivaz et al., 2002). LBPA has also been shown to be necessary for proper dynamics and organization of contents within this inner-membrane system (Kobayashi et al., 1998; Kobayashi et al., 1999). It has recently been proposed that LBPA may influence cholesterol homeostasis beyond the confines of late endosomes and lysosomes, having been shown to be necessary for lipid droplet formation via Wnt signaling within the endoplasmic reticulum, where cholesteryl esters are synthesized (Scott et al., 2015).

To increase cellular LBPA levels in these studies, we used its presumed precursor and structural isomer, phosphatidylglycerol. PG is thought to convert to LBPA along the endo/lysosomal system (Hullin-Matsuda et al., 2007; Poorthuis & Hostetler, 1978), and studies have demonstrated that exogenously administered PG can be converted to LBPA in vivo (Somerharju & Renkonen, 1980), although the anabolic pathway of this conversion remains unknown. The present demonstration that PG supplementation specifically increases the LBPA content of all cells tested supports the hypothesis that PG is a precursor to LBPA. Utilizing PG as a precursor also obviates any potential concern about the LBPA isoform, as the cells presumably generate the physiologically accurate form. Cellular conversion of PG to LBPA has also been demonstrated in mammalian alveolar macrophages (Waite et al., 1987), lymphoblasts (Hullin-Matsuda et al., 2007) and RAW macrophages (Bouvier et al., 2009).

Based on the present results, we propose that the NPC2 hydrophobic ‘knob’ domain inserts into LBPA enriched inner LE/LY membranes, interacting directly with the phospholipid. This, in turn, facilitates rapid transfer of cholesterol from the membranes to NPC2. We speculate, given the ability of NPC2 to promote membrane-membrane interaction, that LBPA and NPC2 are involved in the formation of membrane contact sites which could potentially exist between closely apposed inner LE/LY membranes. Membrane contact sites have been shown to be important in intermembrane lipid transfer (Helle et al., 2013; Holthuis & Levine, 2005; Prinz, 2014), although none have yet been specifically described within the multivesicular LE/LY. The present results show that NPC2 promotes membrane-membrane interaction, and further indicate the ability of NPC2 to bind to LBPA; this interaction may represent one membrane contact point on inner LE/LY membranes. Our previous kinetic studies suggested that NPC2 interacts with other membrane phospholipids as well, and preliminary molecular dynamics simulations indicate that unlike LBPA, where NPC2 interacts at the hydrophobic knob, these interactions occur at NPC2 surface sites other than the knob domain (data not shown); such interactions could represent a second contact point within the inner LE/LY membranes. Interestingly, it has now been demonstrated that NPC2 also binds directly to NPC1 which resides in the limiting LE/LY membrane, and possibly also with LAMP proteins in these same membranes (Li et al., 2016). These could also be potential tether points for the NPC2 and would effectively bring inner LE/LY cholesterol-laden membranes in closer proximity to the limiting LE/LY membrane, which cholesterol must ultimately cross to exit the compartment.

The key finding of the present studies is that LBPA functions in cholesterol efflux, at least in part, by an NPC2-dependent mechanism, as LBPA enrichment in NPC2-deficient fibroblasts was completely ineffective at reversing cholesterol accumulation. This would suggest that an NPC2-dependent, NPC1-indepenedent pathway of cholesterol egress exists, in addition to the ‘hydrophobic handoff’ of cholesterol from NPC2 to NPC1 (Deffieu & Pfeffer, 2011; Infante et al., 2008; Wang et al., 2010). As noted earlier several lines of evidence support NPC2-dependent NPC1-independent cholesterol egress from the LE/LY (Boadu et al., 2012; Goldman & Krise, 2010; Kennedy et al., 2012). Our present demonstration of a critical functional interaction between NPC2 and LBPA also implies that in the presence of intact NPC2, bypassing dysfunctional NPC1 may be achieved via LBPA enrichment. LBPA enrichment may also be effective in NPC2 cases where the mutation is in a residue outside of the hydrophobic knob. For example, a human mutation in D72 has been reported to be disease causing (Biesecker et al., 2009), and we found here that PG supplementation/LBPA enrichment allowed NPC2 deficient cells to be effectively cleared by the D72A protein; supplementation with D72A-NPC2 was entirely ineffective prior to LBPA enrichment.

Currently there is no cure for NPC disease. Pharmacological options are limited, and palliative care remains the standard for treatment of the disease, focusing on increasing the length and quality of life for affected patients. The present study demonstrates that, *in vitro*, NPC disease cells are able to convert administered PG to LBPA. The increase in LBPA occurs concurrently with a decrease in cholesterol accumulation in cells with NPC1 deficiency, which comprises 95% of NPC disease cases. Thus, we propose that cellular LBPA enrichment is worth exploring as a possible therapy. Aerosolized phospholipids such as dipalmitoyl phosphatidylcholine are widely used to enhance pulmonary drug delivery (Duret et al., 2014), and can be adequately nebulized without losing compositional integrity (Schreier et al., 1994). PG itself, administered intranasally, has been used as a therapy by the Voelker group to effectively inhibit respiratory syncytial virus infection (Numata et al., 2010; Numata et al., 2013) and influenza A virus (Numata et al., 2012). Moreover, lipid based colloidal carriers are able to cross the blood brain barrier (BBB) when administered intranasally (Ganesan et al., 2018; Mittal et al., 2014; Patel & Patel, 2017; Tapeinos et al., 2017) and have recently been shown to be efficient drug delivery vehicles in the treatment of intrinsic brain tumors (van Woensel et al., 2013) and neurodegenerative diseases including Alzheimer’s (Agrawal et al., 2018; Tapeinos et al., 2017) and Parkinson’s (Tapeinos et al., 2017; Yang et al., 2016). Given the historical difficulties in treating the neurological effects of NPC disease, the development of a minimally invasive, effective treatment with intrinsic ability to cross the BBB is of interest.

## Materials and Methods

### Orientation of NPC2 in membranes

The Orientation of Proteins in Membranes (OPM) database (http://opm.phar.umich.edu/) was used to predict spatial orientation of NPC2 protein with respect to the hydrophobic core of lipid bilayers. Protein structure of NPC2 (PDB: 1NEP) was searched against the OPM database and the resulting coordinate file and orientation predictions were obtained.

### Sequence alignment and hydrophobicity analysis

Protein sequences for human NPC2 (NCBI Accession: NP_006423.1), rat NPC2 (NCBI Accession: NP_775141.2) mouse NPC2 (NCBI Accession: NP_075898.1), bovine NPC2 (NCBI Accession: NP_776343.1), cat NPC2 (NCBI Accession: XP_003987882.1), chimpanzee NPC2 (NCBI Accession: NP_001009075.1) and the yeast NPC2 (NCBI Accession: KZV12184.1) were aligned with CLUSTAL Omega (Sievers et al., 2011). Protein conservation was scored using a PAM250 scoring matrix, which is extrapolated from comparisons of closely related proteins, similar to the current application (Pearson, 2013). Domain specific conservation of the hydrophobic knob between each NPC2 sequence was analyzed by taking the sum of the conservation scores of each residue from 56 to 64, relative to the human NPC2 sequence. Protein hydrophobicity was scored using the Kyte and Doolittle Amino acid Hydropathicity scale (Kyte & Doolittle, 1982). Domain specific hydrophobicity of the hydrophobic knob of each NPC2 sequence was analyzed by taking the sum of the Kyte and Doolittle Amino acid Hydropathicity score of each residue from 56 to 64. Whole protein hydrophobicity of aligned sequences were analyzed using the ProtScale Tool on the ExPASy server (Gasteiger et al., 2005), based on the Kyte and Doolittle Amino acid Hydropathicity scale (Kyte & Doolittle, 1982).

### Cell Culture

Chinese hamster ovary (CHO) cells transfected with a human NPC2 expression vector (NPC2-800#7) (Liou et al., 2006), kindly provided by Peter Lobel, were maintained in F12-K media (Invitrogen) supplemented with 10% FBS and 1 mg/mL gentamycin. Human WT (GM03652), NPC1 (GM03123), and NPC2 (GM18455) fibroblasts (Coriell Institute, Camden, NJ) were maintained in DMEM media (Invitrogen) supplemented with 15% FBS and 1% penicillin-streptomycin. All cells were at passage 18 or below.

### Generation and Isolation of NPC2 mutants

Point mutations were created with the Stratagene QuikChange Site Directed Mutagenesis Kit (Agilent, Santa Clara, CA), using myc 6xHis-tagged murine NPC2 plasmid, according to the manufacturer’s directions and as described previously (Ko et al., 2003; McCauliff et al., 2015). Plasmid isolation was performed using the PureYield Plasmid Miniprep system (Promega, Madison, WI). Wild type and mutant myc 6xHis-tagged NPC2 proteins were purified from transfected NPC2-800#7 CHO cells, which secrete large amounts of the NPC2 protein into CD CHO media (Invitrogen), using a 10 kDa cutoff flow filtration membrane (Millipore, Bedford, MA) to initially concentrate the media, as previously described (McCauliff et al., 2015). The presence of purified NPC2 was confirmed by SDS-PAGE (Cheruku et al., 2006; Ko et al., 2003; McCauliff et al., 2015); proteins used were > 90% pure by silver staining. Buffer exchange was performed using Sartorius Vivaspin Turbo 4 filters with a 10 kDa cutoff membrane followed by dilution in sodium citrate buffer (in 20 mM sodium citrate, 150 mM NaCl, pH 5.0).

### Cholesterol binding by NPC2

Equilibrium binding constants for cholesterol binding by WT and mutant NPC2 proteins were determined by quenching of tryptophan emission, as previously described (Cheruku et al., 2006; Ko et al., 2003; McCauliff et al., 2015; Xu et al., 2008). Purified proteins were delipidated via acetone precipitation (Liou et al., 2006) and resuspended in sodium citrate buffer. The delipidated NPC2 proteins were incubated with increasing concentrations of cholesterol (>99%) (Sigma Aldrich), in DMSO, for 20 minutes at 25°C and tryptophan emission spectrum were acquired on an SLM fluorimeter (Horiba Jobin Yvon, Edison, NJ). Final DMSO concentration was >1% (v/v). AUC were determined for all spectrum and binding constants were determined by hyperbolic fit of the data using Sigma Plot software (San Jose, CA).

### Membrane vesicle preparation

Small unilamellar vesicles (SUV) were prepared by sonication and ultracentrifugation as previously described (Storch & Kleinfeld, 1986). Large unilamellar vesicles (LUV) were prepared via freeze-thaw cycling and extrusion through a 100nm membrane, as previously described (McCauliff et al., 2015; Xu et al., 2008). The final phospholipid concentration of all vesicles was determined by quantification of inorganic phosphate (Gomori, 1942). Vesicles were maintained above the phase transition temperatures of all constituent lipids. Standard vesicles were composed of 100 mol% egg phosphatidyl choline (EPC) (Avanti Polar Lipids, Alabaster, AL). Where noted, LBPA (Avanti) replaced 5-25 mol% of EPC in SUV and/or LUV preparations, as indicated. All vesicles used for in vitro transfer assays were prepared in sodium citrate buffer, pH 5.0. For incubation with cells, vesicles were composed of 100 mol% PG (Avanti); 100 mol% LBPA or 100 mol% PC (Avanti) and prepared in sterile phosphate-buffered saline, pH 7.4.

### Effect of LBPA on cholesterol transfer by NPC2

As detailed previously (McCauliff et al., 2011; McCauliff et al., 2015; Xu et al., 2008), transfer of cholesterol from WT or mutant NPC2 protein to membranes was monitored by the dequenching of tryptophan fluorescence over time using a stopped-flow mixing chamber interfaced with a Spectrofluormeter SX20 (Applied Photophysics, Leatherhead, UK). Cholesterol transfer rates from 1 μM WT NPC2 to 125 μM EPC membranes containing increasing mol percentages of LBPA (0%, 10%, 20% and 30%) were determined at 25°C. Additionally, to determine the effects of LBPA on the sterol transport properties of mutant NPC2 proteins, transfer of cholesterol from 1 μM WT or mutant NPC2 to 125 μM SUVs composed of either 100% EPC or 25 mol% LBPA/EPC was monitored at 25°C. Instrument settings to ensure the absence of photobleaching were established before each experiment. Data were analyzed with the Pro-Data SX software provided with the Applied Photophysics stopped-flow spectrofluorometer, and the cholesterol transfer rates were obtained by single exponential fitting of the curves, as previously described (McCauliff et al., 2015; Xu et al., 2008).

### Membrane-membrane interaction

Effects of NPC2 on vesicle-vesicle interactions were assessed in two ways, both using light scattering approaches. 200 μM LUVs were mixed with 1 μM WT or mutant NPC2 proteins in a 96-well plate reader, and absorbance at 350 nm monitored every 10 seconds over a period of 30 minutes (Petrusevska et al., 2013). Increases in A350nm (light scattering) are indicative of vesicle-vesicle interaction, the rate of which was determined by a three-parameter hyperbolic fit of the data using Sigma Plot software. Additionally, 750 μM LUVs were mixed with 1 μM WT or mutant NPC2 proteins in a spectrophotometer (Hitachi U-2900, Pleasanton, CA) and A350nm was measured over a period of 60 seconds (Schulz et al., 2009); rates of vesicle-vesicle interaction were obtained by single exponential fitting of the curves.

### Lipid blot analysis of NPC2 interactions

To assess WT NPC2 binding to various LBPA isomers, LBPA Snoopers (Avanti), containing 1 μg spots of pure LBPA isomers, were blocked with tris-buffered saline (TBS) (0.8% NaCl, 20mM Tris-HCl pH 7.4) + 3% BSA (fatty-acid free), followed by a one hour incubation at room temperature with 5 μg of WT NPC2 in TBS + 3% BSA, at a final concentration of 0.5 μg/ml protein. The protein solution was removed and the Snoopers were washed with TBS. NPC2 bound to LBPA isomers was detected by incubating the Snoopers with rabbit polyclonal anti-c-myc-tag antibody (RRID: AB_914457, GenScript, Piscataway, NJ) at a concentration of 0.5μg/ml in TBS + 3% BSA for one hour at room temperature. Following removal of the primary antibody, the strips were washed with TBS and incubated with anti-rabbit IgG HRP-conjugated antibodies (RRID: AB_772206, GE Healthcare, Pittsburgh, PA) at a 1:20,000 dilution in TBS + 3% BSA. After a one-hour incubation with the secondary antibody, the Snoopers were washed with TBS + 0.05% Tween and developed with ECL reagents (GE Healthcare).

For further analysis of NPC2-lipid interaction, Hybond-C membranes (GE-Healthcare) were spotted with either 500 pmol of 18:1 LBPA/BMP (S,R) (Avanti), 18:1 PA (Avanti), 18:1 PG (Avanti), 18:1 PA (Avanti), and Egg PC, or with increasing concentrations of LBPA (125, 250, 375 and 500 pmol) to analyze binding of WT NPC2 to various phospholipid species, or binding of NPC2 mutants to LBPA, respectively.

Following the protocol of Dowler et al (Dowler et al., 2002), each phospholipid was spotted in duplicate and allowed to dry for one hour. Membranes were blocked for 1 hour in blocking buffer containing TBS (50 mM Tris/HCl, pH 7.5, 150 mM NaCl) and 5% (w/v) non-fat dry milk. Membranes were then incubated overnight at 4°C with either WT or mutant NPC2 protein diluted to a final concentration of 1ug/ml in TBS and 3% (w/v) non-fat dry milk. The membranes were then washed at room temperature in TBST (0.1% Tween 20) six times for 5 minutes each, followed by incubation with mouse monoclonal anti-myc antibody (RRID: AB_309938, Millipore) at a 1:2,000 dilution in TBS and 3% (w/v) milk. After washing with TBST, the membrane was then incubated with anti-mouse IgG IRDye-800CW conjugated antibody (LI-COR, Lincoln, NE) at a 1:10,000 dilution in TBS, 0.1% SDS, and 3% (w/v) milk. The membranes were finally washed in TBST 12 times for 5 minutes each at room temperature before acquiring images on the LI-COR Odyssey.

### Homogenous Time Resolved Fluorescence (HTRF)

Assays were performed as described in Fleury, et al. (Fleury et al., 2015), with minor modifications. Reaction mixtures for the interaction assays were prepared in white 384-well polystyrene non-binding surface NBS microplates (Corning, Corning, NY) with a final volume of 20μL per well. Each reaction mix contained 6μL of buffer A (20 mM sodium citrate, 150mM NaCl, 1mM EDTA pH 5.0), 2μL of the recombinant WT or mutant His-tagged NPC2 at a final concentration of 75nM, 2μL of biotinylated lipid solution in buffer A at a final concentration range of 1μM - 58.5nM, 5μL of streptavidin-d2 conjugate (Cisbio Bioassays, Bedford, MA) and 5μL of monoclonal anti-6His-europium cryptate antibody (Cisbio) in detection buffer (20mM of HEPES pH 8.5, 200mM of potassium fluoride, 1% bovine serum albumin). The biotinylated lipids (PS, PA, PG, PC and LPBA; Avanti) were dried under nitrogen and the resultant film was initially reconstituted in EtOH and secondarily in binding buffer at a ratio of 1:10 EtOH:buffer. The final concentration of ethanol in the reaction was 1% (v/v). Following an 18 h incubation of the reaction mixture at room temperature, the fluorescence was measured with an Envision plate reader (Perkin Elmer; λex = 320 nm, λem = 615 and 665 nm; 100 μs delay time). The HTRF ratio value was represented as Ch1/Ch2*10,000 where Ch1 is the energy transfer signal at 665nm, and Ch2 is the europium cryptate antibody signal at 615nm. The negative control wells contained donor and acceptor fluorochromes without NPC2 or biotinylated lipid. The negative control (background) readout was subtracted from all the sample readings.

### Clearance of cellular cholesterol by NPC2 proteins

As detailed previously, a single dose of purified WT or mutant NPC2 protein was added to the media of NPC2 mutant fibroblasts cultured on 8-well tissue culture slides (Nalgene), and allowed to incubate for 3 days. The final concentration of added protein was 0.4 nM. Cells were then fixed and stained with 0.05 mg/mL filipin III (Fisher) and subsequently imaged on a Nikon Eclipse E800 epifluorescence microscope using a DAPI filter set. Filipin stain area was quantified with the accompanying NIS-Elements software (Nikon Inc.) and results were calculated as the ratio of filipin area to total cell area as described previously (McCauliff et al., 2011; McCauliff et al., 2015).

### Cellular LBPA enrichment via PG supplementation

WT, NPC1–, and NPC2 mutant fibroblasts were cultured to confluence in 100mm petri dishes and passaged by trypsinization at a 1:3 ratio. After 24-48 hours, media was removed and replaced by media supplemented with either 30, 100, or 250 μM PG SUVs (Bouvier et al., 2009; Luquain-Costaz et al., 2013), or with 100 μM LBPA or PC SUVs. After 24 hours, cells were collected and 4-6 dishes of the same treatment were pooled. Protein levels were analyzed using the Bradford method (Bradford, 1976). Total cell lipids were extracted from 2 mL of 1 mg/mL protein via the method of Bligh and Dyer (Bligh & Dyer, 1959), resuspended in 200 μL 2:1 chloroform:methanol, and were run on HPTLC plates (EMD Chemicals, Inc) in a solvent of 65:35:5 chloroform:methanol(v/v):30% ammonium hydroxide(v/v) (Akgoc et al., 2015). Lipid spots were quantified by densiometric analysis (ImageJ) from standard curves of authentic standards. TLC results were confirmed with an anti-LBPA antibody (6C4), graciously provided by Jean Gruenberg. Briefly, WT, NPC1– and NPC2 mutant fibroblasts were plated onto 8-well tissue culture slide (BD Falcon) at a density of approximately 6,000 cells/well. Cells were exposed to media containing 100 μM PG SUV, as above, and washed with PBS following the twenty-four hour incubation period. Cells were fixed with 4% paraformaldehyde, rinsed with PBS, and exposed to the 6C4 primary antibody at a 1:1000 dilution in PBS for at least one hour at room temperature or overnight at 4°C. Following removal of the primary antibody, cells were rinsed with PBS, permeabilized with 50 μg/ml saponin in a 10% FBS blocking solution for 5 minutes, rinsed with PBS again and finally exposed to anti-mouse IgG Alexa 488-conjugated secondary antibody (Abcam, Cambridge, MA) at a dilution of 1:200 in PBS for one hour at room temperature. All cells were imaged on a Nikon Eclipse E800 epifluorescence microscope using a FITC filter set to detect the Alexa-488 secondary antibody. Alexa-488 stain area was quantified with the accompanying NIS-Elements software (Nikon Inc) and LBPA accumulation was calculated as the ratio of antibody stain area to cell area.

### Clearance of cholesterol by PG supplementation

WT, NPC1–, and NPC2 mutant fibroblasts were plated onto 8-well tissue culture slides (BD falcon) at a density of approximately 20,000 cells per well and incubated at 37°C, 5% CO_2_ for 24 hours. Culture media was then removed and the cells were incubated with media supplemented with 100 μM PG SUVs for 24 hours. Cells were subsequently fixed and stained with 0.05 mg/mL filipin III and anti-LBPA primary antibodies (6C4), followed by an Alexa 488-conjugated secondary antibody, as described above. In some experiments, NPC2 deficient fibroblasts were secondarily incubated for 24 hours with WT or mutant NPC2 proteins at a final concentration of 0.4 nM, 24 hours after the 24 hour supplementation with PG. Cells were imaged on a Nikon Eclipse E800 epifluorescence microscope using a DAPI filter set to detect filipin and a FITC filter set to detect the Alexa-488 secondary antibody to LBPA. Filipin and antibody stain areas were quantified with the accompanying NIS-Elements software (Nikon Inc.); cholesterol and LBPA accumulation was calculated as the ratio of filipin or antibody area, respectively, to cell area (Delton-Vandenbroucke et al., 2007; McCauliff et al., 2011; McCauliff et al., 2015). To ensure the absence of spectral overlap between the DAPI and FITC filter sets, cells were labeled independently with either filipin or 6C4, and imaged.

### Statistical Analysis

Statistical analysis was performed using OriginPro 2016 (OriginLab Corporation) and SigmaPlot 12.0. Means were compared using Student’s t-test for independent samples or one-way ANOVA where indicated with p < 0.05 considered as significantly different.

## Acknowledgments

This work was supported by funds from the Ara Parseghian Medical Research Foundation (JS, OI), the American Heart Association (predoctoral fellowship 11PRE7330012 to LAM and Grant-in-Aid 14GRNT19990014 to JS), and National Institutes of Health GM 1125866 (JS). The authors would like to thank Joseph Nickels and Hsing-Yin Liu for their assistance with the HTRF assay, Peter Lobel for providing the NPC2-800#7 CHO cells and Jean Gruenberg for generously providing the 6C4 anti-LBPA antibody.

## Author Contributions

The study was conceived and designed by LAM and JS. Experiments were performed by LAM, AL and RL with help from DB and OI. Guidance was provided by PK in protein modeling studies. LAM, AL, RL and JS contributed to writing the manuscript, and all authors contributed to reviewing the manuscript.

